# Purkinje cardiomyocytes of the ventricular conduction system are highly diploid but not regenerative

**DOI:** 10.1101/2022.10.29.514354

**Authors:** Hirofumi Watanabe, Ge Tao, Peiheng Gan, Baylee C. Westbury, Kristie D. Cox, Kelsey Tjen, Ruolan Song, Glenn I. Fishman, Takako Makita, Henry M. Sucov

**Affiliations:** Dept. of Regenerative Medicine and Cell Biology, Medical Univ. of South Carolina; Leon H. Charney Div. of Cardiology, New York Univ. Grossman School of Medicine; Darby Children’s Research Inst., Dept. of Pediatrics, Medical Univ. of South Carolina; Dept. of Medicine, Div. of Cardiology, Medical Univ. of South Carolina

## Abstract

Inefficiency of regeneration underlies many of the pathologies associated with heart injury and disease. Ventricular diploid cardiomyocytes (CMs) are a candidate population that may have enhanced proliferative and regenerative properties [1-3], but subpopulations of diploid CMs and their regenerative capacities are not yet known. Here, using the expression marker Cntn2-GFP and the lineage marker Etv1Cre^ERT2^, we demonstrate that peripheral ventricular conduction CMs (Purkinje CMs) are disproportionately diploid (35%, vs. 4% of bulk ventricular CMs). However, this lineage had no enhanced competence to support regeneration after adult infarction. Furthermore, the CM-specific kinase Tnni3k, which strongly influences bulk ventricular CM ploidy [3] and is also associated with conduction system defects [4], had no influence on the ploidy or organization of the ventricular conduction system. Unlike the bulk diploid CM population, a significant fraction of conduction CMs remain diploid by avoiding neonatal cell cycle activity, likely contributing to these properties.

## Main text

To visualize the ventricular conduction system, we obtained Cntn2-GFP mice, in which GFP is driven by regulatory sequences of the contactin-2 gene in a BAC transgene. Expression of this transgene in the cardiac conduction system has been previously described [5]. We also obtained the Etv1Cre^ERT2^ line, which contains tamoxifen-regulated Cre^ERT2^ knocked into the Etv1 gene [6]. Expression of Etv1-lacZ and Etv1-GFP alleles in the ventricle is limited to the conduction system [7,8]. Because the use of Etv1Cre^ERT2^ for the conduction system has not previously been described, we crossed this allele with Rosa26-tdTomato (R26-tdT), in which tdTomato is conditionally expressed after Cre-mediated recombination, and with Cntn2-GFP. We treated adult mice with tamoxifen to initiate R26-tdT recombination, and visualized GFP and tdT expression patterns on the luminal (endocardial) surfaces of the ventricle (the location of conduction system fibers [9]) by fluorescence. Both reporters showed a near-identical overlapping expression pattern throughout the ventricle (Fig. 1A).

**Fig. 1.**
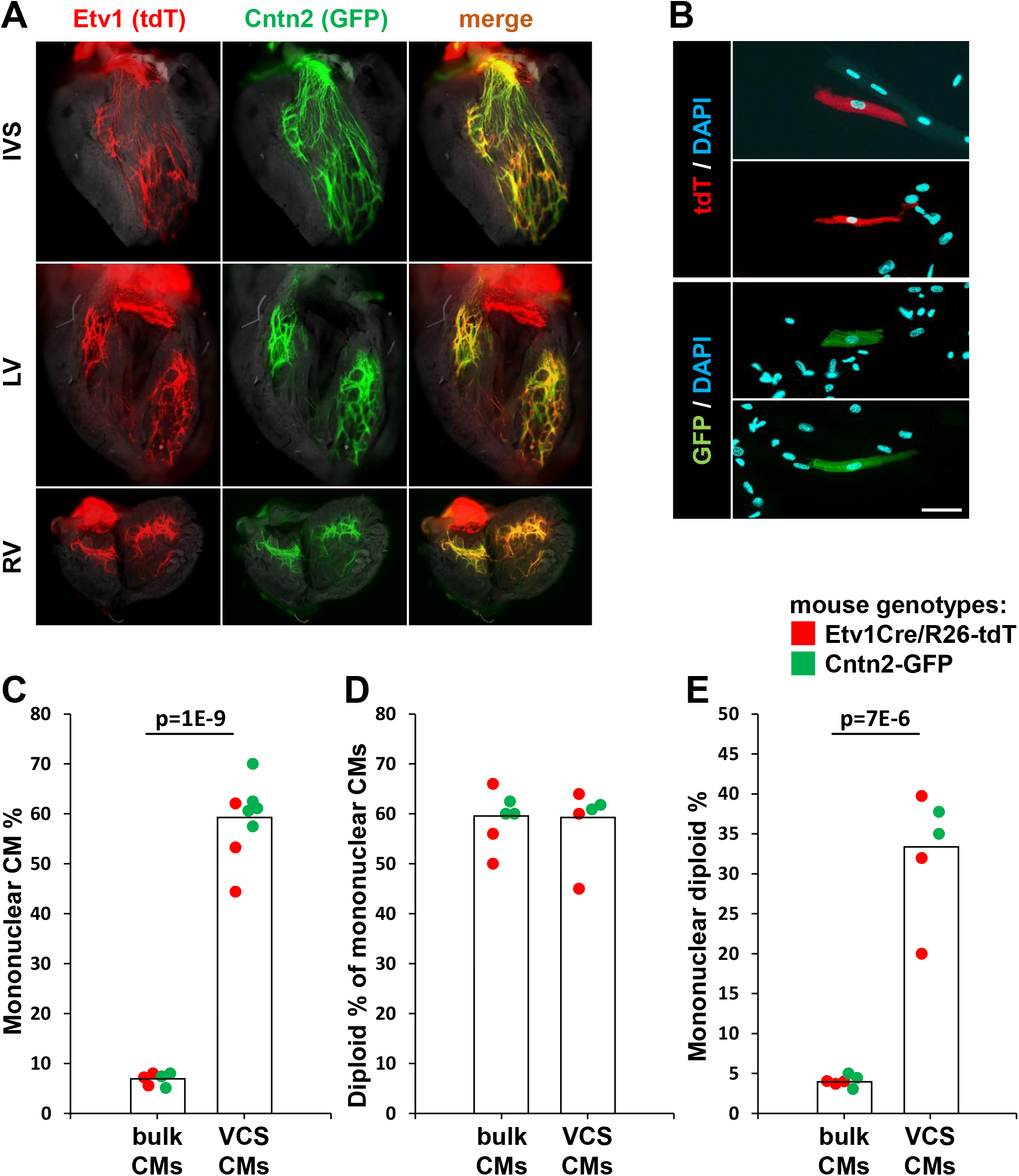
Purkinje CMs are highly diploid. **A**. In situ views of the adult ventricular conduction system labeled by Etv1Cre^ERT2^/R26-tdT and Cntn2-GFP in the same mice. Unfixed hearts were dissected to expose the interventricular septum (IVS), left ventricle (LV) and right ventricle (RV) luminal surfaces. **B**. Examples of isolated mononuclear CMs of diverse morphology expressing either conduction system marker. Scale bar 100µ and the same for all panels. **C-E**. Ploidy analysis of bulk ventricular CMs (not expressing a conduction system marker) or of ventricular conduction system CMs (VCS CMs) expressing tdT (Etv1+) or GFP (Cntn2+) in single cell preparations. Each dot represents one mouse. Because there did not seem to be a substantial distinction in the analysis of the two lines, their data have been combined but color coded to indicate the original source. Three Cntn2-GFP mice were scored for mononuclear CM% but not for nuclear ploidy.

We prepared single cell suspensions of ventricular myocardium from mice carrying either Cntn2-GFP or Etv1Cre^ERT2^/R26-tdT individually. Notably, we utilized only the lower half of the ventricle for this analysis; this preparation is therefore selective for the peripheral component of the ventricular conduction system (Purkinje CMs). With respect to injury and regeneration studies below, this preparation also includes the domains of myocardium that are impacted by left anterior descending coronary artery ligation in experimental infarction. These preparations typically yielded hundreds of thousands of CMs. The recovery of GFP+ or tdT+ CMs was low, varying between 0.02-0.2% of total recovered CMs. This may in part be because our preparations exclude the atrioventricular node and His bundle, where many conduction cells are likely to reside. As previously reported [10], Purkinje cells have a variety of morphologies, and we observed a similar diversity in our preparations, ranging from long and slender to more rectangular or block-like (Fig. 1B).

In these single cell preparations, it was immediately obvious that the labeled cells were disproportionately mononuclear. Mononuclear CMs comprised 7% of the overall ventricular CM population of these mixed-strain mice, a level typical for ventricular populations across many inbred mouse strains [3]. In contrast, over half (59%) of the conduction CMs were mononuclear (Fig. 1C). There did not appear to be an appreciable distinction between Etv1 lineage (tdT+) and Cntn2-GFP+ cells in this or other parameters described below. Because mononuclear CMs may have 2n (n=the haploid genome content) or 4n (or higher) nuclei, we also measured the nuclear ploidy of the mononuclear CM populations. 59% of the mononuclear nonconduction CMs and 58% of the mononuclear conduction CMs were 2n (the remainder being mostly tetraploid, with very few nuclei at higher ploidy level) (Fig. 1D), which is typical for mice as seen in our prior studies [3,11]. Thus, approximately one-third (33%) of the ventricular conduction CMs have one 2n nucleus (1×2n), i.e., are diploid, compared to only 4% of bulk CMs (Fig. 1E).

Virtually all fetal CMs are diploid; CM polyploidy in mouse occurs during the first (and to a lesser extent second) postnatal week when CMs enter cell cycle and replicate their DNA but fail to complete cytokinesis [1,12]. In principle, diploid CMs observed in adult mice may not have entered cell cycle at all (i.e., persisted from embryonic stage) or may have entered and completed cell cycle during this period (i.e., divided). This has not been resolved for the bulk CM population nor for the conduction lineage. To address this, we injected Cntn2-GFP neonatal pups with EdU around noon once per day during the postnatal P2-P5 interval, allowed these mice to age to day 32, and isolated single ventricular cells for analysis. In bulk CMs (nonconduction), early neonatal EdU treatment resulted in labeling of 41±5% (n=3) of CMs overall when examined at P32. The EdU-negative CM group includes cells that either did not have cell cycle activity during the P2-P5 period or cells that were in S-phase during P2-P5 but at a time (e.g., night or early morning) when the level of systemic EdU was too low to result in labeling. Importantly, diploid CMs were EdU+ to an equivalent degree (45±5%) as the overall CM population (41%) and were equally represented in the EdU+ and EdU-populations, in both cases at around 4% (Fig. 2A). These outcomes indicate that diploid CMs in the later postnatal heart did not avoid cell cycle in the earlier neonate in order to remain diploid (if so, they would be disproportionately EdU-), but rather entered and completed cell cycle.

**Fig. 2.**
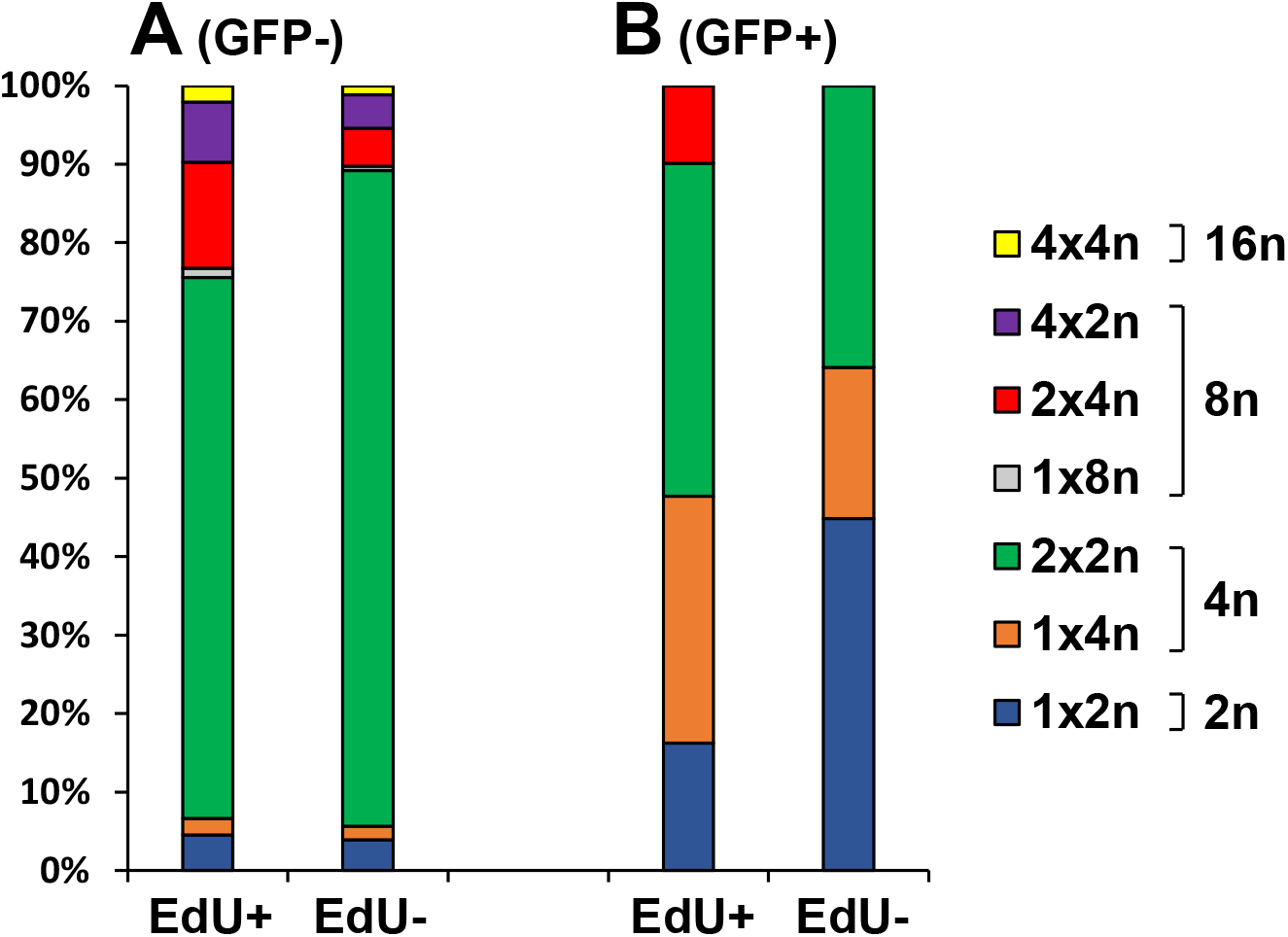
Cell cycle dynamics of bulk (nonconduction) and ventricular conduction (VCS) CMs. Distribution of different ploidy subtypes within the EdU+ and EdU-subsets of: **A**. bulk CMs (GFP-) and of **B**. conduction CMs (GFP+).

Interestingly, EdU+ and EdU- nonconduction CMs differed markedly in the percentage of 4n vs 8n polyploidy (Fig. 2A). A 4n cell (whether 1×4n or 2×2n) is interpreted to having started as a diploid (1×2n) cell, replicated its genome once and then failed to complete karyokinesis or cytokinesis. An 8n cell is interpreted to have arisen from a 4n cell that replicated its genome again and then again failed in cytokinesis. A higher fraction of the EdU+ bulk CM group was 8n (22% vs. 10%, p=0.006), and reciprocally, a higher fraction of the EdU-population was 4n (85% vs. 71%, p=0.009). This indicates that CMs that became EdU labeled during the P2-P5 period were more likely to enter cell cycle again later, thereby reaching a higher ploidy level. A higher fraction of the EdU- group conducted only one round of cell cycle activity and remained at a lower ploidy level.

When the GFP+ CMs from the same mice were analyzed, a much different profile emerged. Here, a smaller percent of GFP+ CMs overall were EdU+ (29±3%; compared to 41% of GFP-CMs), and only 13±2% of the diploid CMs were EdU+ (compared to 45% for bulk CMs; p=0.004). Consequently, a much higher percent of the EdU-subgroup was diploid (45% vs. 16%, p=0.02; Fig. 2B, 1×2n group). This indicates that the conduction CM lineage disproportionately avoids cell cycle entry during the P2-P5 period, and implies that a substantial fraction of these cells persisted from the embryo into the postnatal period as diploid cells.

Tnni3k status influences CM ploidy [3,11], specifically by impacting the mononuclear percentage with no impact on nuclear ploidy. Because the conduction system population is so highly diploid, we addressed whether Tnni3k has a specific influence on the conduction system. To do so, we separately crossed the Etv1Cre^ERT2^/R26-tdT and Cntn2-GFP conduction system reporters into the Tnni3k mutant background [3]. By whole mount fluorescence, no prominent difference in the organization of the ventricular conduction system was evident between control and mutant adult mouse hearts (Fig. 3A). In ventricular single cell preparations, absence of Tnni3k did not change the mononuclear percentage of the conduction cells when data from both reporters were combined (Fig. 3B). There may be a slight impact of Tnni3k mutation on the Etv1Cre lineage, although this was not statistically significant. The absence of a more prominent effect of Tnni3k mutation on the conduction lineage is likely for two reasons. First, Tnni3k influences the percentage of neonatal CMs that complete cytokinesis after cell cycle entry, whereas a high fraction of the conduction lineage does not enter cell cycle in the first place (Fig. 2). Second, because this population is already so highly diploid in control hearts, any impact of Tnni3k mutation to further enhance cell cycle completion would be difficult to detect.

**Fig. 3.**
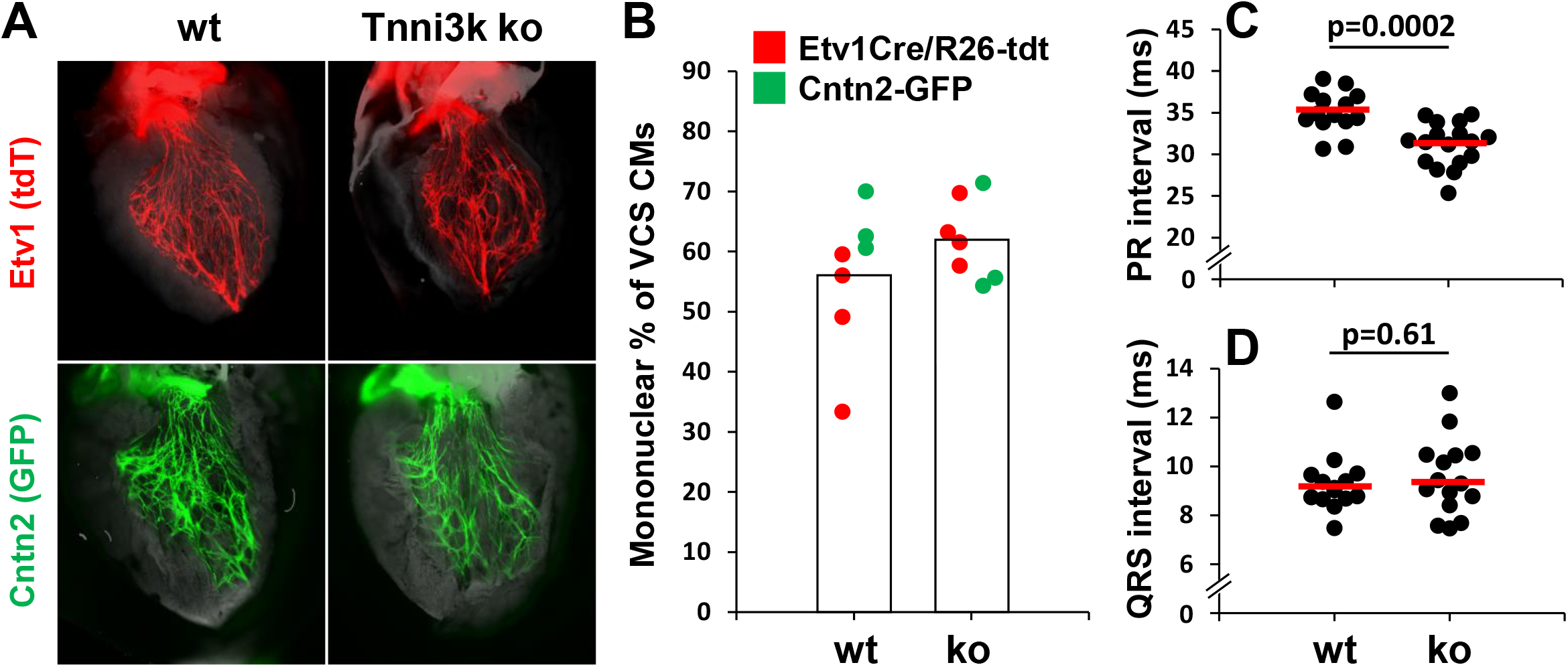
Tnni3k does not influence the Purkinje cell lineage. **A**. In situ views of the interventricular septa of unfixed adult hearts from wild-type and Tnni3k mutant mice carrying either conduction system label. **B**. Minimal change in mononuclear percentage within the conduction CM subset in Tnni3k wild-type (wt) vs. mutant (ko). Data for the two reporters were combined but color coded as in Fig. 1; there may be a trend to a difference specifically within the Etv1Cre lineage although this was not statistically significant (p=0.07). **C-D**. PR and QRS intervals measured by ECG in C57BL/6-inbred wild-type and Tnni3k mutant adult mice.

Some genetic manipulations in mice have been shown to alter CM polyploidy and to cause arrhythmias [13,14], although it is not known if either is causal to the other. In human pedigrees, several different mutations in the TNNI3K gene have been associated with a variety of arrhythmias and related conduction system abnormalities [4]. In mice, absence of Tnni3k has been shown to cause a shortening of the PR interval although not of additional conduction phenotypes [15]. We performed lead II surface ECG under isoflurane anesthesia to address if there is a functional impact of Tnni3k mutation on the ventricular conduction system. There are two differences in our analysis compared to the previous mouse study [15]. First, here we used an engineered complete loss-of-function allele [3] whereas the past study addressed the natural mutant allele; because the natural allele is a 5’ splice site mutant at the intronic +5 position [15], it may support some degree of correct splicing (i.e., may be a partial hypomorph rather than a full null). Second, all of the mice evaluated here were on an inbred C57BL/6J strain background, and strain background could have a prominent impact on conduction phenotypes. As in the previous analysis, we observed a similar shortening of the PR interval in mutant mice (Fig. 3C). More germane to this study, the QRS interval, which is reflective of the functionality of the ventricular Purkinje system [9], was unchanged in mutant mice (Fig. 3D), as also reported in the prior study [15]. Heart rate was equivalent in the two groups (wild-type: 611±37 bpm; mutant: 621±29 bpm, p=0.38). We did not observe other ventricular phenotypes that might be reflective of a Purkinje function defect, such as ventricular tachycardia. In sum, we conclude that Tnni3k mutation does not alter the spatial distribution, ploidy composition, or baseline function of the ventricular conduction lineage in an appreciable manner. Equally important is that conduction defects (as in the human TNNI3K studies) are unlikely to arise from the influence of Tnni3k on ventricular conduction CM ploidy, as this lineage is already so highly diploid and this is at most only marginally changed based on Tnni3k status.

We next considered the possibility that the Purkinje lineage might be preferentially competent to support proliferation, particularly after adult myocardial infarction. This was suggested by several observations. First, diploid CMs are a candidate CM subtype that may have elevated regenerative ability [2,3], and as reported above the conduction lineage is highly diploid.

Second, lineage tracing in the injured adult heart revealed that proliferative CMs are preferentially located on the endocardial side of the ventricle [16], which is where conduction CMs are also located. Third, the relatively small number of Purkinje CMs is compatible with the low level of CM cell cycle activity in adult hearts as measured by static markers or nucleotide incorporation.

For this analysis, adult Etv1Cre^ERT2^/R26-tdT mice were treated with tamoxifen and then injured by permanent LAD coronary artery ligation (Fig. 4A). We inserted an Alzet minipump at the time of infarction to release EdU, removed this after 14 days, and after 2 more days isolated hearts or heart cells for analysis. Histology of the intestine isolated at the same time confirmed the effectiveness of this EdU treatment protocol (Fig. 4D). Sections of the recovered hearts revealed prominent infarctions across the left ventricular anterior wall and extending into the ventricular septum (Fig. 4B). Staining in the infarcted region and border zone revealed EdU labeling in numerous nonmyocytes and in occasional CMs (Fig. 4B). Rare instances of EdU+tdT+ double labeled cells were also noted.

**Fig. 4.**
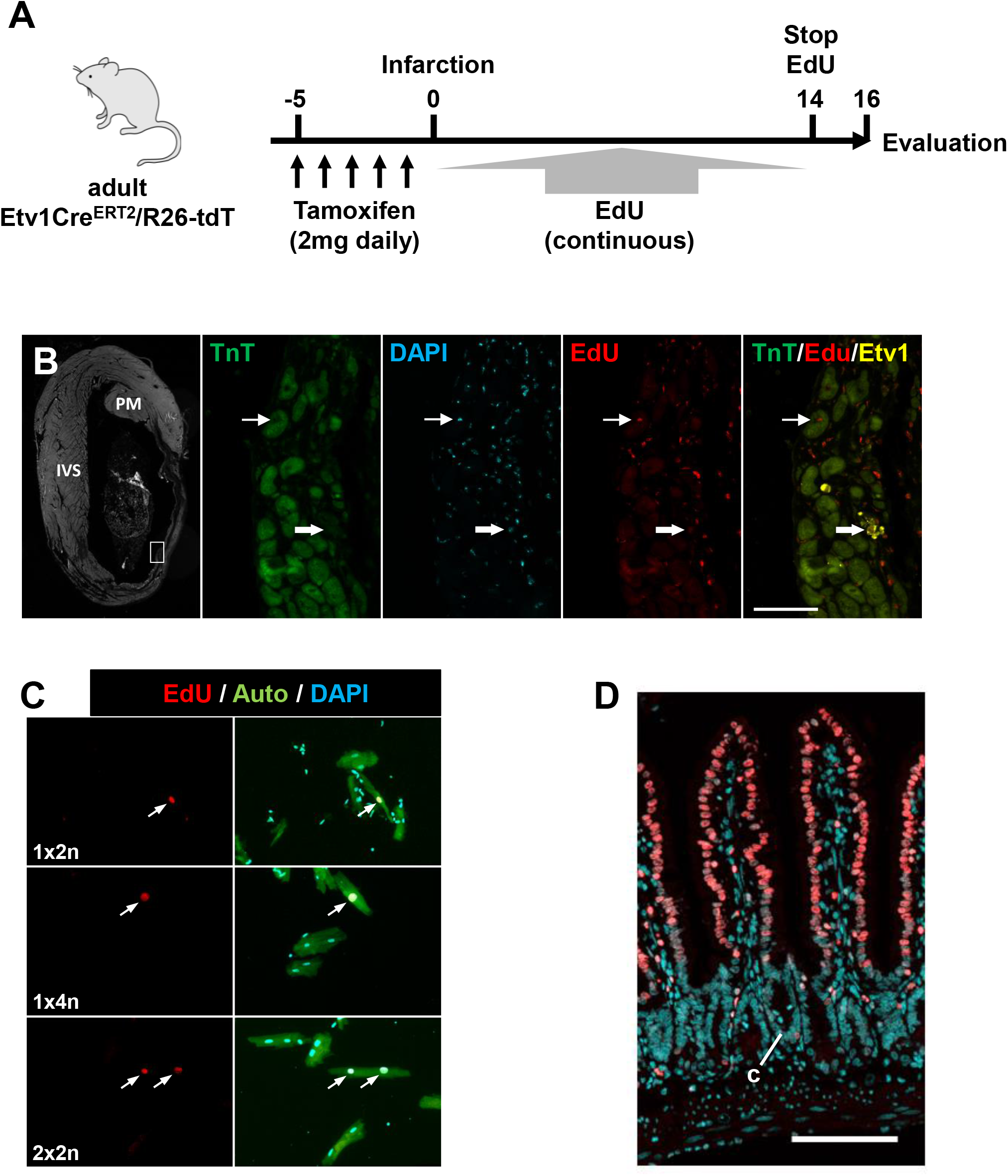
The Purkinje lineage does not contribute to heart regeneration. **A**. Schematic of the protocol for infarction and EdU labeling. **B**. Histology of infarcted heart and cell cycle activity within the Etv1 lineage. The small box in the first panel indicates the location in the infarction border zone where the higher magnification views were obtained. PM, papillary muscle; IVS, interventricular septum. The higher magnification panels show an example, indicated by the thick arrow, of an Etv1-lineage labeled CM that is EdU+. Nearby is an EdU+ CM that is not a conduction CM (indicated by the thin arrow). Scale bar 100µ. **C**. Examples of EdU+ CMs in single cell preparations isolated from infarcted hearts. 1×2n, mononuclear diploid; 1×4n, mononuclear tetraploid; 2×2n, binucleated with two diploid nuclei. Auto, autofluorescence in the green channel (specific for CMs). **D**. Histology of the small intestine of a mouse taken 16 days after infarction and 2 days after removal of the EdU pump. Because of the 2-day period without label, the crypts (c) are unlabeled whereas the villi are labeled from midway to tip, indicating the persistence of EdU throughout the labeling period. Scale bar 100µ.

We prepared ventricular single cell suspensions from infarcted hearts (Fig. 4C), recovering an average of 459 ± 125 EdU+ CMs per heart (n=3 hearts). Importantly, in these preparations no EdU+ CMs were derived from the Etv1 lineage. The conduction lineage is a small fraction of the total CM pool, so the absence of recovered EdU+tdT+ cells does not exclude that conduction cells engage in cell cycle after injury. However, these results do indicate that the ventricular conduction lineage does not have a unique ability to do so, despite its high degree of diploidy.

As noted above, relative to the bulk CM population a much higher proportion of the conduction lineage in the newborn heart avoids cell cycle activity and thereby persists as diploid; this same tendency to not enter cell cycle may also occur in the injured adult heart. These observations do not contradict the premise that diploid CMs are more highly regenerative than polyploid CMs, as the Purkinje lineage is only one subset of the total diploid CM population. A separate subpopulation, yet to be defined, may account for the enhanced regeneration associated with diploid CMs.

## Methods

### Mice

The BAC-transgenic Cntn2-GFP line was originally sourced from MMRRC (stock 029940-UCD) and obtained from Glenn Fishman (NYU). The Etv1Cre^ERT2^ knock-in line was obtained directly from JAX (strain 013048). The conditional R26-tdTomato line was obtained from JAX (strain 007914). All three alleles were outcrossed to a variety of strain backgrounds before this study began. To induce recombination in Etv1Cre^ERT2^/R26-tdT mice, tamoxifen (T5648, Sigma-Aldrich; 100 mg/kg body weight in corn oil) was administered by i.p. injection once per day for 5 days, with subsequent analysis no earlier than 48hr after the last injection. Tnni3k knockout mice were previously described by us [3]. All mice used for experimental purposes in this study were adults, generally of 3-6 months of age, and both sexes were used with no sex-based differences noted.

### Whole-mount fluorescence

Using fresh isolated hearts, an incision was first made along the long axis of the anterior surface of the left ventricle to expose the ventricular septal surface and the left ventricular free wall. The right ventricle was then exposed through an incision along the long axis of the anterior surface of the right ventricle. Each interior surface was photographed by fluorescence microscopy (Keyence BZ-X810) using the automatic image stitching function.

### Ventricular single cell preparations

As previously described [3,17], hearts were removed from anesthetized heparin-treated mice, immersed in ice-cold Kruftbrühe (KB) solution (70mM potassium aspartate, 40mM KCl, 15mM KH_2_PO_4_, 10mM glucose, 10mM taurine, 0.5mM EGTA, 10mM sodium pyruvate, 10mM HEPES, 5mM BDM, 0.5% BSA) to stop contraction, then quickly attached to a Langendorff perfusion rig. Hearts were perfused with calcium-free Tyrode’s solution (120mM NaCl, 4mM KCl, 0.33mM NaH_2_PO_4_, 1mM MgCl_2_, 10mM HEPES, 11mM glucose, 20mM taurine, 20mM BDM) and then digested with 50mg collagenase type II (1 mg/ml) in calcium-free Tyrode’s solution. Ventricular tissue alone (the lower half of the ventricle) was manually removed using forceps and then triturated by gentle pipetting in KB solution to make a single cell suspension. This was filtered by gravity through a 250μ nylon mesh and then fixed in 2% paraformaldehyde (PFA) in PBS at room temperature for 15 min. In EdU labeling studies, EdU was visualized in cell suspensions using a Click-iT EdU AlexaFluor 594 Imaging Kit (C10339, Thermo Fisher Scientific). When used, cell preparations were immunostained with mouse anti-cardiac troponin T antibody [1C11] 1:500 (ab8295, Abcam), subsequently with goat anti-mouse IgG (H+L) cross-adsorbed secondary antibody AlexaFluor 488 conjugate 1:1000 (A11001, Thermo Fisher Scientific) to define CMs; in other cases CMs were identified by their natural autofluorescence in the green channel. Nuclei were stained with 5μg/ml Hoechst 33342 (H3570, Thermo Fisher Scientific) or 5μg/ml DAPI for 5min. Cells were washed in PBS then applied to microscope slides and mounted with ProLong Gold Antifade Mountant (P36930, Thermo Fisher Scientific).

### Nuclear number and ploidy

Numbers of nuclei per cardiomyocyte and their ploidy were quantified using photographs taken at a uniform setting for all cell preparations in an experiment with a Leica DFC3000G camera in full frame mode (1296 × 966 pixels) on an Olympus BX41 microscope (20x objective), or with a Keyence BZ-X810 with a built-in camera (1920 × 1440 pixels). Mononuclear percentage was defined by counting at least 100 cells per slide from at least 5 slides per heart preparation. Nuclear ploidy of cardiomyocytes was measured based on DAPI or Hoechst fluorescence intensity in images, calculated using ImageJ and normalized to the median intensity value of binucleated CM nuclei (defined as diploid).

### ECG analysis

Mice were anesthetized by isoflurane vapor inhalation administered via a Kent Scientific SomnoSuite anesthesia system with a MouseSTAT pulse oximeter and heart rate module and taped in a supine position to a warming pad. Needle electrodes were inserted subcutaneously in lead II configuration. ECG data were acquired using an AD Instruments PowerLab 8/35 with Animal Bio Amp, a Bridge Amp, and BP Transducer/Cable kit. After 5-10 min without recording, data were captured using LabChart Pro Data Acquisition & Analysis software. Periods of uniform waveform free from spurious signals were identified and PR and QRS measurements made by averaging 10 consecutive beats.

### Infarction and analysis

Tamoxifen was administered to Etv1Cre^ERT2^/R26-tdT mice by i.p. injection once per day for 5 days, and the day after the last dose, mice were subjected to left anterior descending artery (LAD) ligation. After LAD ligation, an Alzet pump (model 1002, 14-day capacity; Durect Corp.) containing 50mg/ml EdU (A10044, Thermo Fisher Scientific; dissolved in DMSO then diluted 1:1 with saline) was implanted subcutaneously under the back skin. 14 days later, the Alzet pumps were removed under light anesthesia, and 2 days later the mice were euthanized and hearts removed for analysis.

### EdU analysis of neonatal mice

Cntn2-GFP neonatal mice were subcutaneously injected with 200μg of EdU in 50μl (consisting of 2μl of a stock solution in DMSO diluted with 48μl PBS) once daily from P2 to P5. Isolation of cardiomyocytes was performed at P32, with EdU and GFP staining as above. 60 nonoverlapping images were taken at 20x magnification and the number of nuclei and ploidy of all cardiomyocytes evaluated as above.

### Histology

Freshly isolated hearts were immersed in ice cold KB buffer to arrest in diastole and fixed in 4% paraformaldehyde (PFA) in PBS at 4°C overnight with gentle agitation on a rocker. Hearts were dehydrated through increasing ethanol concentrations then embedded in paraffin, 5μ sections were obtained with a Leica RM 2135 microtome. Antigen retrieval was performed in a microwave oven for 25min in a pressure cooker using citrate antigen retrieval buffer (10mM citrate, 0.05% Tween 20, pH 6.0). For EdU detection, Click-iT EdU AlexaFluor 594 Imaging Kit (C10339, Thermo Fisher Scientific) was used following the product instructions. After subsequent blocking with 5% blotting grade milk (1706404, Bio-Rad) for 2hr, heart tissue sections were incubated at 4°C overnight with primary antibodies: mouse anti-cardiac troponin T antibody [1C11] 1:500 (ab8295; Abcam); rabbit anti-RFP at 1:200 (200-301-379; Rockland). On the following day, heart tissue sections were incubated at room temperature for 2 hr with secondary antibodies: goat anti-mouse IgG (H+L) cross-adsorbed secondary antibody AlexaFluor 488 conjugate 1:500 (A11001, Thermo Fisher Scientific); goat anti-rabbit IgG (H+L) cross-adsorbed secondary antibody AlexaFluor 647 conjugate 1:500 (A21244, Thermo Fisher Scientific). Intestines were removed freshly and formed into Swiss rolls [18] and fixed overnight in 10% formalin. The intestines were embedded in paraffin and cut into 5µ sections in the same manner as above. After deparaffinization, Edu detection was performed as above. Subsequently, nuclear staining was performed with 5μg/ml Hoechst 33342 (H3570, Thermo Fisher Scientific) for 15min.

